# Genome-wide chromatin binding of transcriptional corepressor Topless-related 1 in *Arabidopsis*

**DOI:** 10.1101/2020.04.25.060855

**Authors:** Thomas Griebel, Dmitry Lapin, Barbara Kracher, Lorenzo Concia, Moussa Benhamed, Jane E. Parker

**Author notes:** Corresponding author – Jane E. Parker. Evotec (München) GmbH, Munich, Germany (Barbara Kracher). First co-authors (Thomas Griebel, Dmitry Lapin).

## Abstract

Timely and specific regulation of gene expression is critical for plant responses to environmental and developmental cues. Transcriptional coregulators have emerged as important factors in gene expression control, although they lack DNA-binding domains and the mechanisms by which they are recruited to and function at the chromatin are poorly understood. Plant Topless-related 1 (TPR1), belonging to a family of transcriptional corepressors found across eukaryotes, contributes to immunity signaling in *Arabidopsis thaliana* and wild tobacco. We performed chromatin immunoprecipitation and sequencing (ChIP-seq) on an *Arabidopsis TPR1-GFP* expressing transgenic line to characterize genome-wide TPR1-chromatin associations. The analysis revealed ∼1400 genes bound by TPR1, with the majority of binding sites located at gene upstream regions. Among the TPR1 bound genes, we find not only regulators of immunity but also genes controlling growth and development. To support further analysis of TPR1-chromatin complexes and other transcriptional corepressors in plants, we provide two ways to access the processed ChIP-seq data and enable their broader use by the research community.

## Introduction

Transcriptional corepressor proteins regulate gene expression despite lacking recognizable DNA binding domains and are proposed to rely on transcription factors (TFs) for their activity at specific chromatin regions (Chen and Courey, 2000; Causier et al., 2012). Homologs and functional analogs of the corepressors Groucho (Gro) in *Drosophila melanogaster* and Thymidine uptake 1 (Tup1) in *Saccharomyces cerevisiae* are present across eukaryotes (Chen and Courey, 2000; Liu and Karmarkar, 2008). In *Arabidopsis thaliana* (hereafter *Arabidopsis*), the Gro/Tup1 family comprises 13 members divided into two groups: one group includes homologs of Topless (TPL) and Wuschel Interacting Proteins (WSIPs) and the second group, Leunig (LUG) and its homologs (Liu and Karmarkar, 2008). Full-length Gro/Tup1 family proteins carry a C-terminal WD40-repeat domain and diverse N-terminal domains mediating intermolecular interactions (Chen and Courey, 2000; Liu and Karmarkar, 2008; Ke et al., 2015; Martin-Arevalillo et al., 2017). Such interactions with DNA binding TFs, histone deacetylases (HDAC), mediator components and repressor-domain proteins help to confer and maintain a repressed chromatin state (Chen and Courey, 2000; Gonzalez et al., 2007; Liu and Karmarkar, 2008; Pauwels et al., 2010; Thireault et al., 2015).

*Arabidopsis* TPL and its four homologs (TPR1 to 4; <u>TP</u>L-related) emerged as regulators of diverse biological processes, including development (Long et al., 2002; Long et al., 2006) and signaling by the hormones auxin and jasmonates (Szemenyei et al., 2008; Pauwels et al., 2010; Acosta et al., 2013). Evidence points to TPR1 also participating in immunity signaling (Zhu et al., 2010; Niu et al., 2019). Indeed, immunity caused by a mutated, autoactive immune receptor-like protein Suppressor of NPR1, Constitutive 1 (SNC1) in *Arabidopsis* and pathogen-triggered receptor N in *Nicotiana benthamiana* require a functional *TPR1* gene (Zhu et al., 2010; Zhang et al., 2019).

To date, no chromatin binding profile for plant Gro/Tup1 corepressors is available, which limits our understanding of their corepressor activities. Here, we describe a genome-wide TPR1 chromatin binding profile obtained via chromatin immunoprecipitation coupled to sequencing (ChIP-seq) in leaves of autoimmune *Arabidopsis* transgenic plants expressing *pTPR1:TPR1-GFP* (Zhu et al., 2010). To simplify access to the data, we provide input-normalized TPR1 ChIP-seq data which can be directly visualized in the IGV browser. We also make available here a set of R scripts together with necessary input files that will help researchers obtain TPR1 binding profiles (metaplots) for their gene sets of interest.

## Results and Discussion

To study TPR1-chromatin associations *in planta*, we used an *Arabidopsis* Col-0 line expressing *TPR1-GFP* under control of a 2 kb upstream sequence (*pTPR1:TPR1-GFP* Col-0; referred to as *TPR1-GFP*) (Zhu et al., 2010). This line is autoimmune and displays up-regulated defenses, pathogen resistance and growth retardation (Zhu et al.,2010). We performed ChIP-seq experiments with α-GFP antibodies in three independent biological replicates using leaves of six-week-old *TPR1-GFP* plants. The *pTPR1:TPR1-HA* #3 Col-0 line (hereafter *TPR1-HA*) served as a negative control as it is also autoimmune and displays enhanced immunity to the adapted downy mildew pathogen (Zhu et al., 2010). We applied a double linear DNA amplification (LinDA) protocol (Shankaranarayanan et al., 2011) before preparing sequencing libraries. LinDA was instrumental for ChIP-seq based profiling of WRKY TF/chromatin interactions in *Arabidopsis* (Liu et al., 2015; Birkenbihl et al., 2017; Birkenbihl et al., 2018).

In total, we identified 1531 peaks on the *Arabidopsis* genome with a TPR1-GFP signal enriched over input and *TPR1-HA* negative control samples (Supplemental Data 1). These loci were associated with 1441 gene annotations (TPR1 bound genes) (Supplemental Data 2). Most peaks were located in gene promoter regions (47%, 1.0 kb upstream of transcription start site (TSS); Fig. 1a). A further 28% of peaks were found in annotated intergenic regions and 20% of peaks mapped to transcription termination sites (TTS). A small fraction of peaks (<5%) corresponded to intronic and exonic regions (Fig. 1a). A metaprofile of the input-normalized *TPR1-GFP* ChIP-seq signal across 1441 TPR1 bound genes confirmed its primary association site upstream of TSS (Fig. 1b, upper panel). A heatmap showing read densities for individual TPR1 bound genes reinforced this conclusion and further indicated that a fraction of TPR1 bound genes has TPR1 binding at both TSS and TTS (Fig. 1b, lower panel). The association of transcription regulators with both TSS and TTS was also observed in ChIP-seq analyses of WRKY (Birkenbihl et al., 2017) and Myelocytomatosis 2 (MYC2) (Wang et al., 2019) TFs. Thus, we concluded that the TPR1-chromatin association profile generally resembles binding profiles of TFs.

**Fig. 1.**
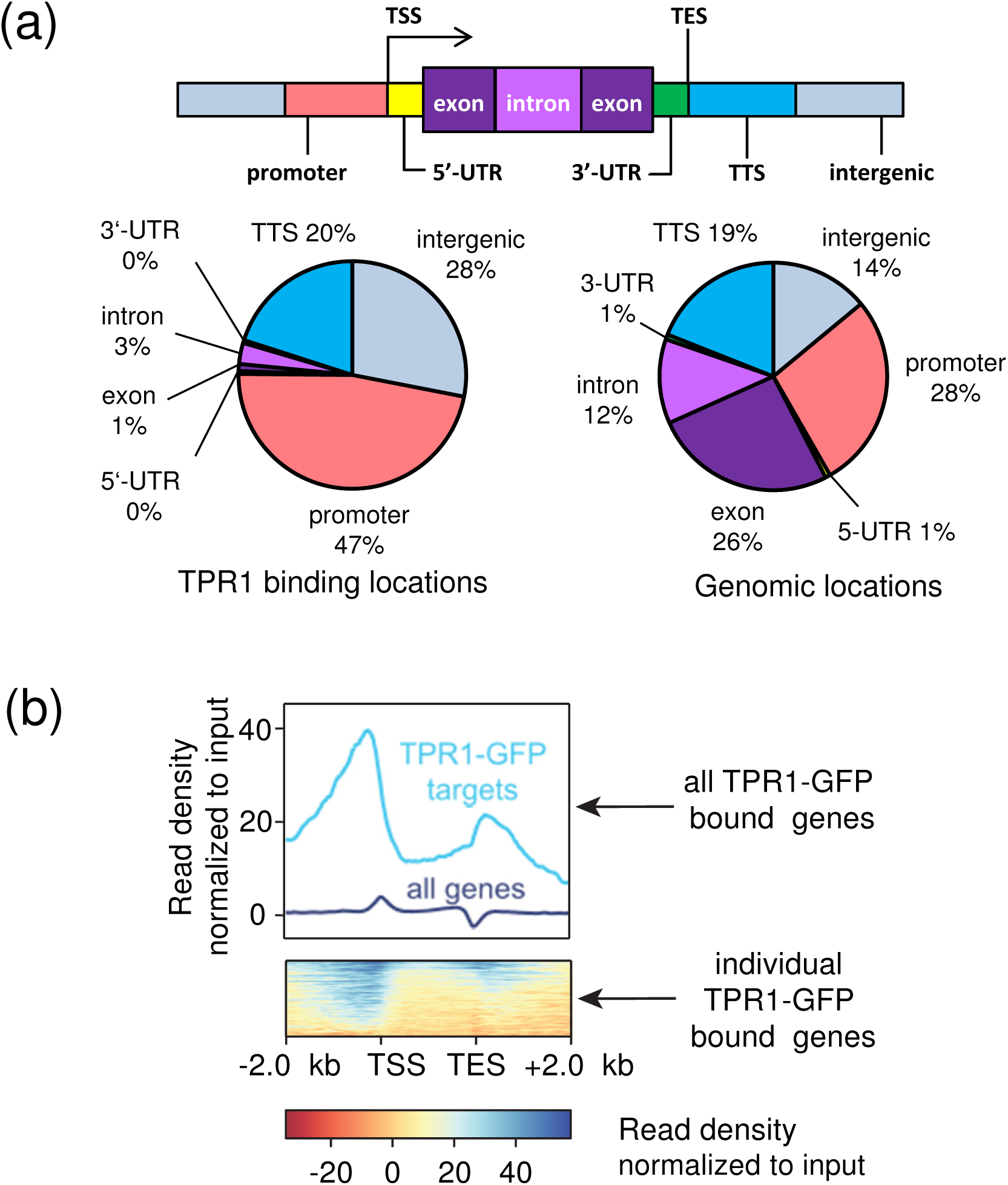
ChIP-seq reveals 1441 genomic TPR1 binding sites mainly located in gene promoter regions upstream of transcription start sites. (a) Pie chart showing the distribution of TPR1 bound genomic regions (ChIP-seq peaks) over *Arabidopsis* genome partitions (left) and the distribution of these partitions in the genome (right). Promoters are defined as 1000 bp regions upstream of the transcription start sites (TSS), and transcription termination sites (TTS) as the 1000 bp region downstream of the 3-UTR. Genomic regions in between a TTS and the promoter of the next gene are considered as intergenic. (b) Metaprofile of 1441 TPR1 bound genes and all *Arabidopsis* genes (control). The TPR1-GFP read density was normalized to the input (upper panel). Heatmap showing the input-normalized read density of the 1441 individual TPR1-GFP bound genes. Read density is shown over gene bodies including a ±2 kb flanking region. (a) and (b) Figures are based on the pooled samples from three independent ChIP-seq replicates.

The TPR1 bound genes showed an enrichment of gene ontology (GO) terms associated with abiotic and biotic stress responses, shoot developmental processes, and responses to plant hormones such as jasmonates, abscisic and salicylic (SA) acids (Fig. 2; Supplementary Data 3). Examples of TPR1 bound genes in development are several auxin-induced genes as *SAUR14* (*Small Auxin Upregulated RNA 14*; Fig. 3). Our ChIP-seq analysis also confirmed that TPR1 binds the promoter of *Defense No Death 1*(*DND1;* Fig. 3) (Zhu et al., 2010). Notably, promoters of several genes induced by pathogens and involved in SA metabolism emerged as TPR1 bound (Figure 3). These genes include *Isochorismate Synthase 1* (*ICS1*) and *avrPphB Susceptible 3* (*PBS3*) important for isochorismate derived SA accumulation (Wildermuth et al., 2001; Rekhter et al., 2019; Torrens-Spence et al., 2019), as well as *Downy Mildew Resistant 6* (*DMR6*) encoding an SA deactivating 5-hydroxylase (Zhang et al., 2017). These observations suggest that TPR1 is directly involved in host transcriptional reprogramming of immune responses. How TPR1 binding to promoters influences gene transcription and the chromatin environment needs to be clarified. It is remarkable that Gro/Tup1 proteins such as *Arabidopsis* Leunig homolog are also associated with gene expression activation (Chen and Courey, 2000; You et al., 2019).

**Fig. 2.**
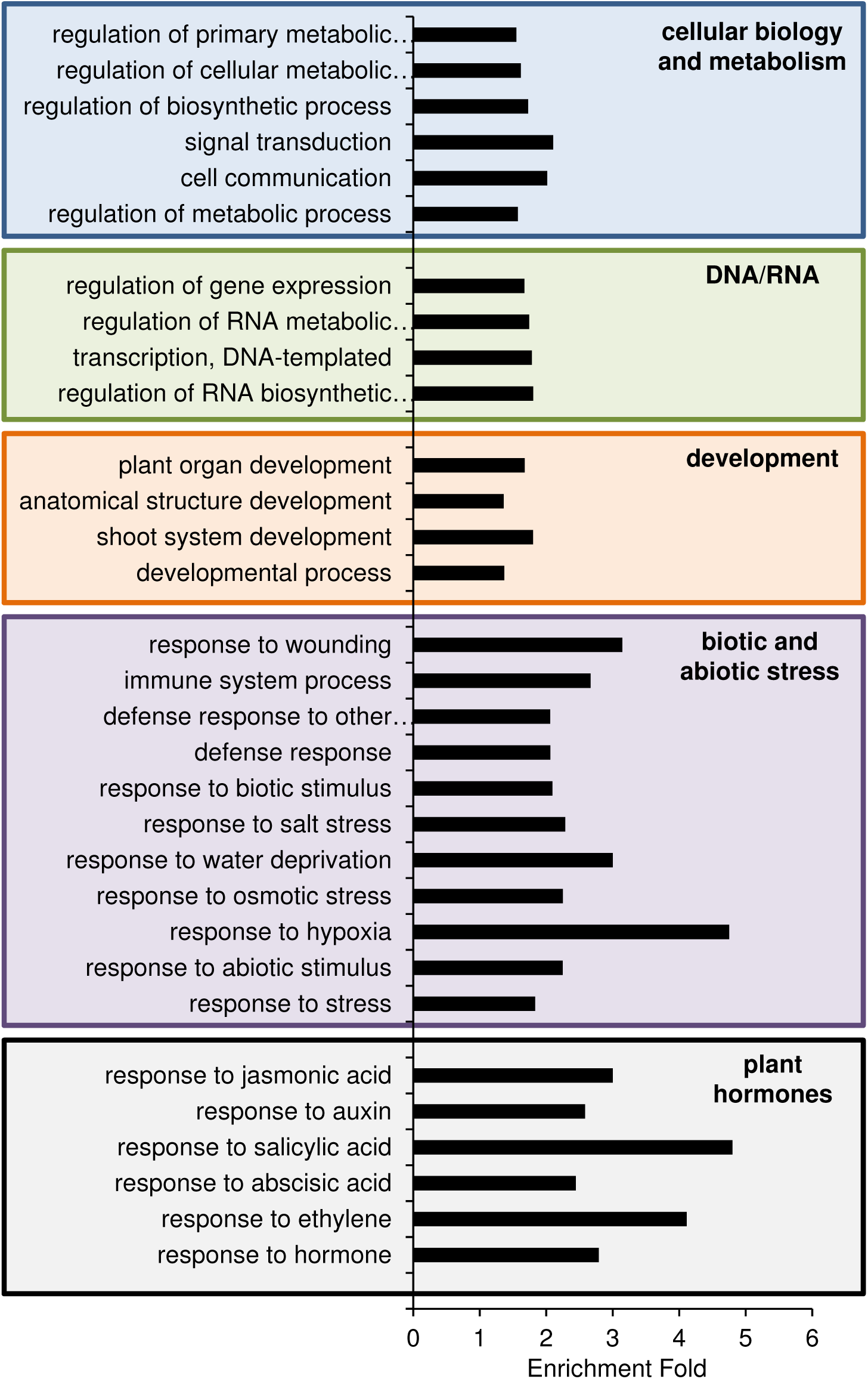
Fold enrichment for selected representative gene ontology (GO) terms from the analysis of 1441 TPR1 bound genes. Selection of total 114 GO terms of biological processes identified for the TPR1 binding targets (corrected p < 0.001; according to default settings using https://go.princeton.edu/cgi-bin/GOTermFinder. The enrichment is calculated to relative to the whole genome GO annotation. The complete list of GO terms is provided in Supplemental Data 3.

**Fig. 3.**
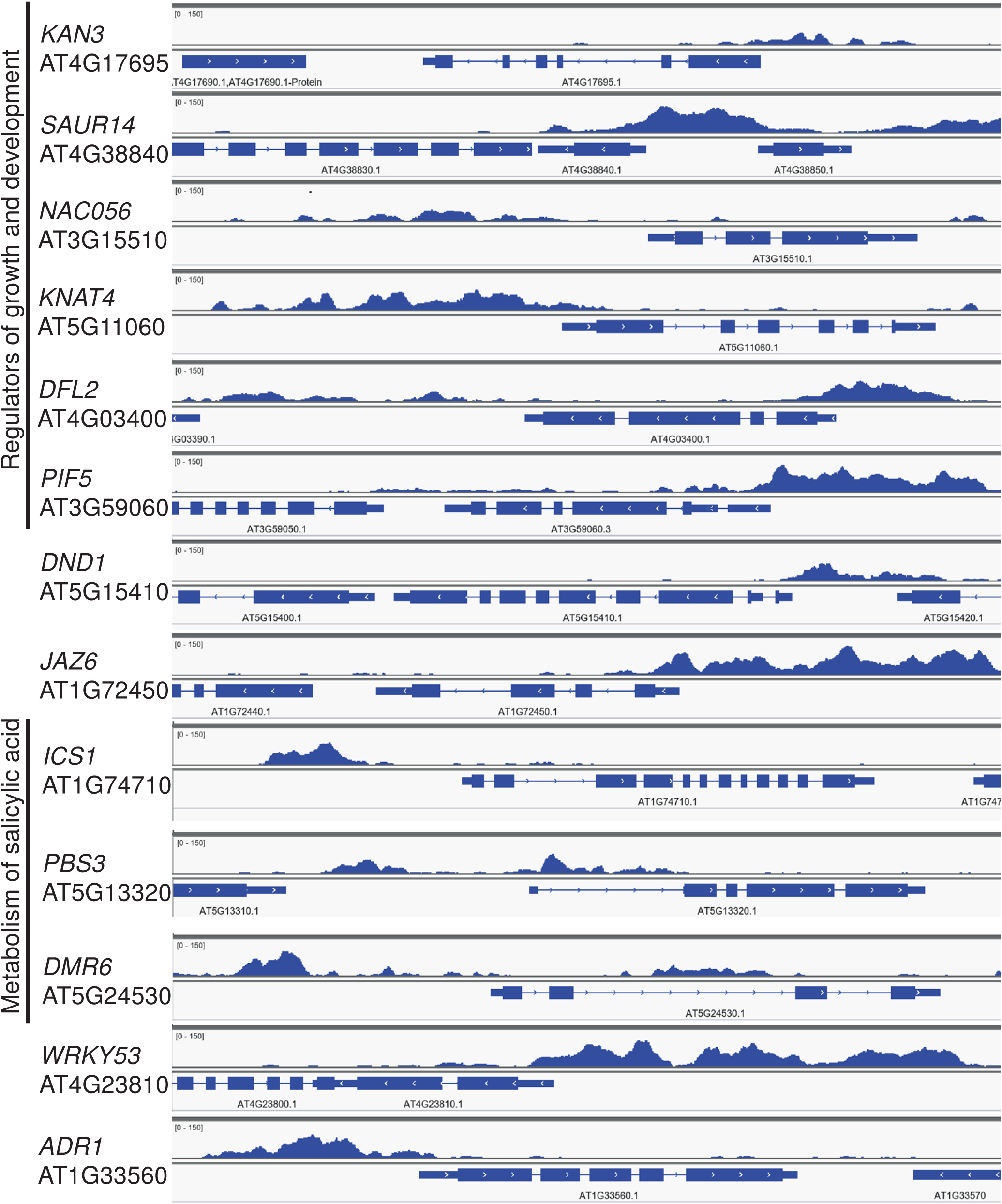
Input-normalized ChIP-seq profiles of TPR1-GFP at selected TPR1 bound genes. The screenshots were taken from Integrative Genome Viewer (IGV) using the input bigwig file “substr_TPR1_WT_bs1.bw” deposited at GEO (accession number 149316).

To support functional and molecular studies of TPR1 and other Gro/Tup1 proteins in *Arabidopsis*, we provide here not only the raw read data but also processed ChIP-seq analysis files that can be used by the community without having to process read data. First, researchers could directly examine TPR1-GFP binding to genes of interest by feeding the bigwig file with input-normalized read density (see Material and Methods for download instructions) to the desktop installation of Integrative Genome Viewer (IGV) (Thorvaldsdóttir et al., 2013). Second, the metaplots will be informative for initial genetic and molecular investigations. To help researchers test for enrichment of TPR1 at the groups of genes of their interests, we release two R scripts to use in combination with the mapped reads (bam files) (see Material and Methods). With this material, metaplots can be obtained on a personal computer (∼8 Gb RAM) with little (if any) coding.

In summary, this study describes the first genome-wide profile of *Arabidopsis* TPR1 associations with nuclear chromatin. The information will likely contribute to our understanding of transcriptional reprogramming in plant immunity responses and guide further studies of other Gro/Tup1 protein functions in Arabidopsis and related species.

## Material and Methods

### Plant materials and growth conditions

Stable transgenic lines *pTPR1:TPR1-GFP* Col-0 (*TPR-GFP*) and *pTPR1:TPR1-HA* #3 Col-0 (*TPR1-HA*) were described previously (Zhu et al., 2010). Plants were grown on soil with a daily 10 h light period (150-200 µE/m^2^s) at 22 °C and 60 % relative humidity. Plant leaf material was harvested after 6 weeks of growth.

### Chromatin immunoprecipitation assays and sequencing (ChIP-seq)

ChIP was performed on 2 g leaf tissues of 6-week-old plants according to Gendrel et al. (2005) with modifications (Birkenbihl et al., 2012). Leaf samples were cross-linked twice by infiltration with 1% formaldehyde under a vacuum for 2x 7.5 minutes. Nuclei were disrupted by sonication 10 times for 30 s each and 30 s of cooling in a Bioruptor (Diagenode). IPs were performed using α-GFP (Ab6556, Abcam) antibodies and Protein A agarose beads (Sigma). After elution of immune complexes, cross-linking was reversed with an overnight incubation at 65°C in 200 mM NaCl followed by proteinase K digestion. DNA was extracted with phenol-chloroform and precipitated with ethanol in the presence of 0.3 M sodium acetate (pH=5.2) and 10 µg/mL glycogen (overnight at -20°C). After centrifugation, DNA pellets were washed with 70% ethanol, air-dried and resuspended in water before PCR. MiniElute PCR Purification Kit (Qiagen, Germany) was used for cleaning ChIP DNA and two rounds of *in vitro* transcription by T7 RNA polymerase followed according to a linear DNA amplification (LinDA) protocol (Shankaranarayanan et al., 2011). Sequencing libraries were barcoded and generated using Ovation® Ultralow DR Multiplex system (NuGEN Technologies Inc.) including a 16-cycles PCR amplification. Libraries were size-selected (200-350 bp) with an agarose gel extraction using the MinElute Gel Extraction Kit (Qiagen, Germany). Sequencing was performed on Illumina HiSeq2500 by the Max Planck-Genome-centre Cologne, Germany (http://mpgc.mpipz.mpg.de/home/) resulting in about 10-12 million 100 bp single-end reads per sample (Supplemental Data 4). ChIP-seq data are deposited in the National Center for Biotechnology Information Gene Expression Omnibus (GEO) database with accession number 149316.

### ChIP-seq data analysis

Before mapping, remaining LinDA adapters and low quality sequences were removed from the sequencing data using a two-step procedure. Here, first Bpm and t7-Bpm sites were trimmed from the 5′ end using cutadapt (version 1.2.1) with options–e 0.2, -n 2 and–m 36 (otherwise default settings were used), and subsequently poly-A and poly-T tails and low quality ends were trimmed and reads with overall low quality or with less than 36 bases remaining after trimming were removed using PRINSEQ lite (version 0.20.2) (Schmieder and Edwards, 2011) with options–trim_qual_right/left 20, trim_tail_right/left 3 –min_len 36, -min_qual_mean 25. After the preprocessing steps, remaining high quality reads were mapped to the *A. thaliana* reference genome TAIR10 (http://www.arabidopsis.org) using Bowtie (version 2.0.5; default settings) (Langmead and Salzberg, 2012). For subsequent removal of non-uniquely mapping reads, the alignment output was filtered for mapping quality using samtools (version 0.1.18) (Li et al., 2009) view with option -q 10.

To identify genomic DNA regions (‘peak regions’) enriched in sequencing reads in the ChIP sample compared to the input control and *TPR1-HA* negative control, the peak calling algorithm of the QuEST program (version 2.4) (Valouev et al., 2008) was applied using the TF mode (option ‘2’), with permissive parameter settings for peak calling (option ‘3’). Each of three biological replicates was first analyzed separately. To obtain more exact peak locations for consistent peaks, mapped reads of the three replicates were additionally pooled and peaks called for the pooled samples. Further analyses were performed on peak regions identified from the pooled samples. To annotate peak locations with respect to annotated gene features in TAIR10 the annotatePeaks.pl function from the Homer suite (Heinz et al., 2010) was used with default settings.

### Generation of metaplots and the bigwig file for Integrative Genome Viewer (IGV)

A metaplot was generated with the suite deepTools3.0 (Ramírez et al., 2014) using functions bamCompare (--operation subtract, default ‘readCount’ scaling), computeMatrix (scale-regions) and plotHeatmap. Bed files for gene regions were prepared based on TAIR10 annotation (TAIR10_GFF3_genes_transposons, 11-07-2019). Raw sequence reads were trimmed with cutadapt (version 1.9.1, -e 0.2 -n 2 - m 30) to remove overrepresented sequences detected with fastqc (version 0.11.9) (Andrews et al., 2010) and then mapped to TAIR10 using bowtie2 (version 2.2.8). The resulting BAM files from individual biological replicates were deduplicated, filtered for low-quality mapping (-q 10 in view function in samtools version 1.9) and finally merged with samtools v1.9 (Li et al., 2009; Li, 2011). The bigwig file provided for visualization in IGV (Thorvaldsdóttir et al., 2013) was prepared with bamCompare (--operation subtract --binSize 1) using merged BAM files for input and TPR1-GFP samples. This bigwig file (∼240 Mb, “substr_TPR1_WT_bs1.bw”) is available for download on GEO (149316) and through the respective Max Planck Digital Library collection (MPDL; https://edmond.mpdl.mpg.de/imeji/collection/U6N5zIOIWgjjMZCu).

### Preparation of metaplots for gene sets of interest in R

To provide an alternative way to access ChIP-seq data, we prepared two R scripts that generate bed files for a user-provided set of genes (“01_Preparation_BED_files.R”) and then draw metaplots (“02_Drawing_metaplots_TPR1.R”) using the R package metagene (v 2.18.0). These scripts and a guide are available via the Github repository (https://github.com/rittersporn/Griebel_TPR1_bioRxiv_Apr2020). BAM and the respective BAI files were submitted to the Edmond collection at MPDL (https://edmond.mpdl.mpg.de/imeji/collection/U6N5zIOIWgjjMZCu) and should be used as inputs. The scripts were tested on Windows 10 (PC, RAM 8 Gb, Intel Core i5-6600, 3.3 GHz, RStudio v1.1.463, R version 3.6.1 (2019-07-05)) and MacOS Catalina 10.15.2 (MacBook Pro, RAM 8 Gb, Dual Core Intel Core i5, 2.7 GHz, RStudio 1.2.5042, R version 3.6.3 (2020-02-29)). R version ≥ 3.6 is required.

## List of supplemental data files

Supplemental data file 1: List of TPR1-GFP peaks and their annotation

Supplemental data file 2: List of genes annotated as bound by TPR1-GFP

Supplemental data file 3: GO term analysis for TPR1 bound genes

Supplemental data file 4: Overview of raw and mapped read numbers for each sample

## Acknowledgement

This work was supported by the Max Planck Society and Deutsche Forschungsgemeinschaft (DFG) (grants CRC1403 B08 to J.E.P., D.L. and CRC670 TP19 to J.E.P., T.G.). We thank Yuelin Zhang for providing the *pTPR1:TPR1-GFP* and *pTPR1:TPR1-HA* lines and Rainer Birkenbihl for helpful advice on ChIP methodology and the LinDA protocol. We also thank the Max Planck-Genome-centre Cologne (http://mpgc.mpipz.mpg.de/home/) for sequencing of the samples in this study.

## Author contributions

TG and JEP designed the experiment; TG performed the experiment; TG, DL, BK, LC, MB, JEP analyzed the data; DL prepared the Github repository and materials to access processed ChIP-seq data; TG, DL and JEP wrote the manuscript with input from all authors.

